# Oligo-FISH Validates Genome Assemblies and Delivers the Most Precise Karyotype for *Lens* Mill. Species

**DOI:** 10.1101/2025.06.02.657502

**Authors:** Alex Junior Aparecido Silvestrini, Larissa Ramsay, Eric Bishop von Wettberg, Kirstin E. Bett

## Abstract

Chromosome structural rearrangements play a significant role in karyotype evolution and speciation. These rearrangements pose challenges for precise karyotyping, leading to asymmetric chromosomes and complicating the assembly of a genus pan-genome for crops and their wild relatives. *Lens culinaris*, an important cool-season legume primarily cultivated in India and Canada, is the cultivated species among six wild relatives. All seven species of *Lens* face significant challenges due to chromosomal rearrangements, ranging from introgression issues to difficulties in developing a precise karyotype and advancing genomic studies. Using the gene synteny analysis between the cultivated *Lens* species and six wild relatives, we developed cross-species oligo-FISH (Fluorescent *in situ* hybridization) probes aiming to further attest to the genome assembly and synteny analysis of *Lens* species. Roots of seven *Lens* spp. accessions were harvested and used for chromosome spread preparations. Those slides were then used for Oligo-FISH experiments, where the DNA present in the slides was denatured, and a set of red and green oligo probes was hybridized to the chromosomes. Pictures were taken using a fluorescence microscope. The combination of both oligo sets/probes resulted in a distinct pattern for each *Lens* spp. chromosome, allowing the inference of the most precise karyotype to date for six *Lens* species. The number of oligo probe signals reflects the species’ phylogenetic proximity, while the distribution of those signals changed drastically within the same gene pool. The karyotyping of *Lens* confirmed the proper assignment of chromosomes in the genome assemblies and validated the rearrangements detected in the synteny analysis. Differences in the assembly probe prediction and the oligo-FISH results were used to improve the assemblies. The results attest to a higher sequence-level similarity among the closest related species despite the occurrence of several structural changes among them. The oligo-FISH probes can be used in conjunction with plant genome assembly projects, supporting the delivery of a precise representation of their physical chromosomes.

## Introduction

The genus *Lens* Mill. belongs to the family Fabaceae and legume tribe Vicieae, and is comprised of seven taxa – the cultivated *Lens culinaris* and six wild relatives (Alo et al., 2011; Guerra-García et al., 2021; Wojciechowski et al., 2004). All *Lens* species are diploid, have 2n = 14 chromosomes (Galasso, 2003) and are classified into four gene pools depending on the feasibility of crossing with the cultivated species. Lentil is an important pulse crop behind only chickpea and pea in cool-season legume global production (Ritche et al., 2023). The use of wild relatives for lentil breeding is a strategy for positive allele transfer (Cao et al., 2024; Kumar et al., 2018; Marshall et al., 2024; Singh et al., 2018; Tullu et al., 2011, 2013; Vargas et al., 2024) despite the issues that arise when trying to cross species from different gene pools. Most of the crossing issues can be associated with the different chromosome arrangements among those species (Davies et al., 2007; Laskar et al., 2019; Ramsay et al., 2021; Silvestrini et al., Unpublished; Singh et al., 2018). *Lens* species are also of interest for chromosome evolution studies due to the ploidy level and chromosome number stability, coupled with the occurrence of several chromosome changes.

Chromosome rearrangements have been observed in *Lens* species using chromosome staining at F1 lines meiosis (Ladizinsky, 1979; Ladizinsky et al., 1984, 1985), genetic linkage maps (Cao et al. 2024; Gujaria-Verma et al., 2014; Sharpe et al., 2013) and more recently, genome assembly synteny analysis (Silvestrini et al., 2025). *Lens* species karyotype was previously built based on chromosome measurements and fluorescent in situ hybridization (FISH) using repetitive DNA sequences as marks (Balyan et al., 2002; Gaffarzade et al., 2007; Galasso, 2003; Galasso et al., 2001). Although very informative, none of those works confirmed the occurrence of the numerous rearrangements observed when comparing genome assemblies (Ramsay et al., 2021; Cao et al., 2024; Silvestrini et al., Unpublished). There is still no consensus regarding the karyotype of the seven *Lens* species, with authors proposing different karyotype formulas and different numbers and locations of ribosomal DNA sequences (Jha, 2022).

Karyotype studies can be a powerful tool to characterize chromosome rearrangements *in situ*, providing physical evidence of the chromosomal changes and overcoming synteny analysis limitations (Liu et al., 2018). Karyotypes built based on chromosome measurements can provide indirect evidence of rearrangements between different species (Schubert et al., 1991), whereas fluorescent *in situ* hybridization (FISH) marks can provide direct evidence of those changes. The recently developed Oligo-FISH technique can be used as a powerful resource to physically validate both the assignment of scaffolds to chromosomes and synteny analysis in genome assembly projects. The technique is based on the development of conserved oligo probes from genome assembly data covering specific genomic regions. The probes can be used in cross-species FISH experiments, allowing the inference of chromosome evolution, chromosomal compatibility as well as building strong karyotypes for the analyzed species (Braz, do Vale Martins, et al., 2020a; Braz et al., 2018; Braz, Yu, et al., 2020; Jiang, 2019).

We developed an oligo-FISH barcode system based on genome synteny analysis among seven *Lens* species. We strategically defined genomic regions in the cultivated species (*L. culinaris*) that would cover the maximum number of rearrangements between the cultivated and wild relatives. Our system confirms the rearrangements detected through the synteny analysis, as well as allows the validation of the genome assemblies at the chromosomal level and the quality assessment of both the genome assembly and the synteny analysis. The results are a strong karyotype for six *Lens* species, which takes into account the rearrangements that shaped chromosome evolution across the genus and are very useful to assess chromosomal compatibility in plant breeding programs without the need of genotype specific genome assemblies.

## Material and Methods

### Plant Materials and Genome assembly

Genome assemblies of the seven *Lens* species were retrieved from https://knowpulse.usask.ca/ (Table 1).

**Table 1.**
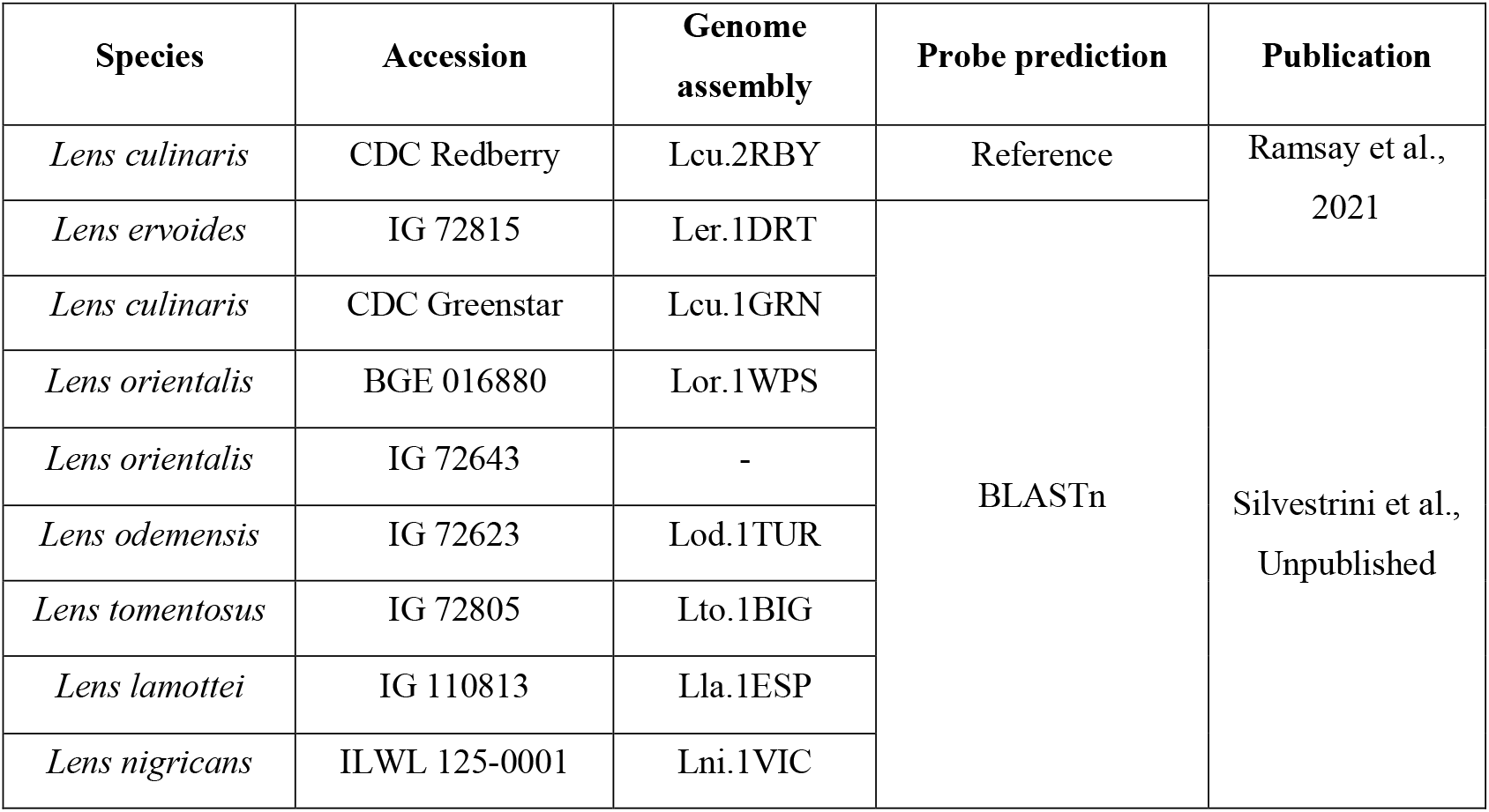
Species and genomes used for Oligo-FISH. Probe prediction method from the reference genome and the BLASTN in the wild *Lens* genomes.

### Oligo probe development and analysis

Genomic coordinates spanning 16 chromosomal regions (Table 2) of the reference genome *L. culinaris* CDC Redberry (Lcu.2RBY) were chosen based on the predicted genome rearrangements from mapping of Lcu.2RBY against Ler.1DRT, Lor.1WPS, Lto.1BIG, Lod.1TUR, Lla.1ESP using bedtools intersect (Quinlan & Hall, 2010). Lcu.2RBY genomic coordinates and the full assembly were sent to Arbor Biosciences (Ann Arbor, MI, USA) for probe development. The repetitive sequences in the Lcu.2RBY genome were filtered out, and the remaining sequences were divided into oligos of approximately 45 nt with a step size of 5 nt. These oligos were processed to remove duplicated oligos and oligos located within the centromeric regions to avoid centromere-specific sequences and possible unfiltered repeats. Two final oligo pools were synthesized – Lcu.2RBY_Gpool labelled with Alexa488 for green signal and Lcu.2RBY_Rpool labelled with ROX for red signal.

**Table 2.**
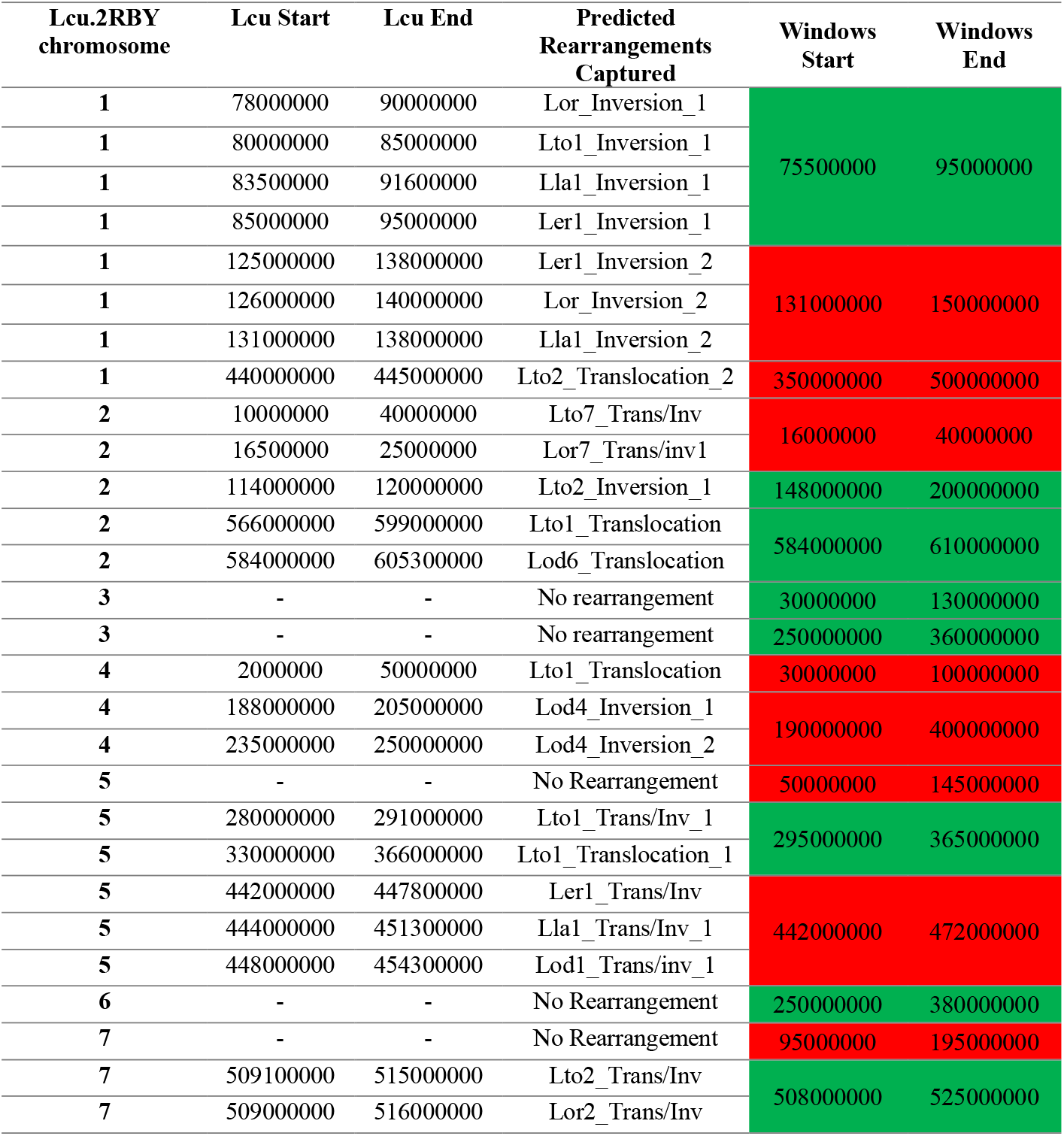
Final genomic windows selection for oligo probes development based on the Lcu.2RBY genome assembly. WS: Windows Start. WE: Windows End. Abbreviations follow the same pattern as Sup Table 1.

The density of oligos in each probe for all other *Lens* species assemblies was determined through BLASTN analysis. Oligo probe annotation was conducted by comparing gene-annotated genomic positions in Lcu.2RBY with oligo probe positions using bedtools intersect (Quinlan & Hall, 2010). All figures were generated using R package ggplot2 (Wickham, 2016), Synvisio (Bandi & Gutwin, 2020), JBrowse 2 (Cain et al., 2022), or IGV (Robinson et al., 2011).

### Fluorescent *in situ* Hybridization using Oligo Probes

Seeds for each of the lines (Sup. Table 1) were germinated in the dark overnight, kept growing for 3 days and treated with 1.18 mM Hydroxyurea (HU) solution for 18 hours to synchronize the cell cycle. Seedlings were left in media without HU for cell cycle recovery, with times varying depending on species (Sup. Table 1), and subsequently treated with 15 μM Oryzalin, a mitotic spindle blocker, to induce metaphase arrest for 4 hours. Roots were then fixed in Carnoy solution (3 parts ethanol: 1 part acetic acid) and stored at 10 ºC until further use. Root tips were subjected to cell wall digestion using an enzyme mixture of one part cellulase and two parts pectinase at 37 ºC with times described in Sup. Table 1, depending on the species. After enzymatic digestion, slides were prepared following the protocol described by Aliyeva-Schnorr et al. (2015). Slides displaying optimal chromosome spread were selected for Oligo-FISH (Fluorescent *in situ* hybridization) using the oligo probes. The Oligo-FISH protocol established by Braz, Yu, et al. (2020) was used with minor adjustments. Briefly, selected slides were pre-fixed with 4% formaldehyde for 15 minutes, followed by three washes with 2x SSC solution. Chromosomes on the slides were denatured at 85 ºC for 2 minutes. Subsequently, slides were incubated overnight with a hybridization mix containing 1 uL of each oligo probe at 37 ºC. Stringency washes with 2x SSC (Saline-sodium citrate) were performed post-hybridization, and images were captured using a ZEISS Axio Imager Z1 Apotome fluorescent microscope with excitation filters for DAPI, Alexa488 and ROX. At least five metaphases per species were analyzed to determine the distribution of probes on chromosomes. Captured images were processed for RGB channel merge using Adobe Photoshop 24.1.1. Images of *in situ* chromosomes were compared with the BLASTn expected pattern from both sets of oligo probes for each analyzed species. Any deviations in the observed patterns were documented and correlated with the number of oligos in each probe at specific genomic regions.

## Results

### Oligo-FISH probe design strategy

The genome rearrangements described in the synteny analysis of Ramsay et al., 2021 and Silvestrini et al., Unpublished, were the basis for the oligo probe design strategy. Synteny comparisons between the Lcu.2RBY genome assembly and five other wild genomes – Lor.1WPS, Lto.1BIG, Lod.1TUR, Lla.1ESP and Ler.1DRT (Table 1) generated precise Lcu.2RBY rearrangement coordinates (Sup. Table 2). The rearrangements mapped to Lcu.2RBY leveraged coordinates that overlap the maximum number of rearrangements against the wild genomes (genomic windows), thus maximizing coverage of the Lcu.2RBY oligo probes in regions of rearrangements (Sup. Table 3).

The final coordinates for each Lcu.2RBY genomic window are presented in Table 2 and Figure 1, colour-coded for each oligo pool (red and green). Each genomic window has at least 17 Mbp coverage, providing sufficient space for 3,375 oligos, each 45 nt in length. Additionally, each window is located at least 15 Mbp from chromosome ends, ensuring a safe distance from telomeric repeats, and is at least 28 Mbp away from adjacent windows. This ensures adequate resolution to differentiate between oligo probes, as proposed for Maize, which has larger chromosomes (Braz, do Vale Martins, et al., 2020b). The genomic windows were sent to Arbor Biosciences for oligo probe development and synthesis.

**Figure 1.**
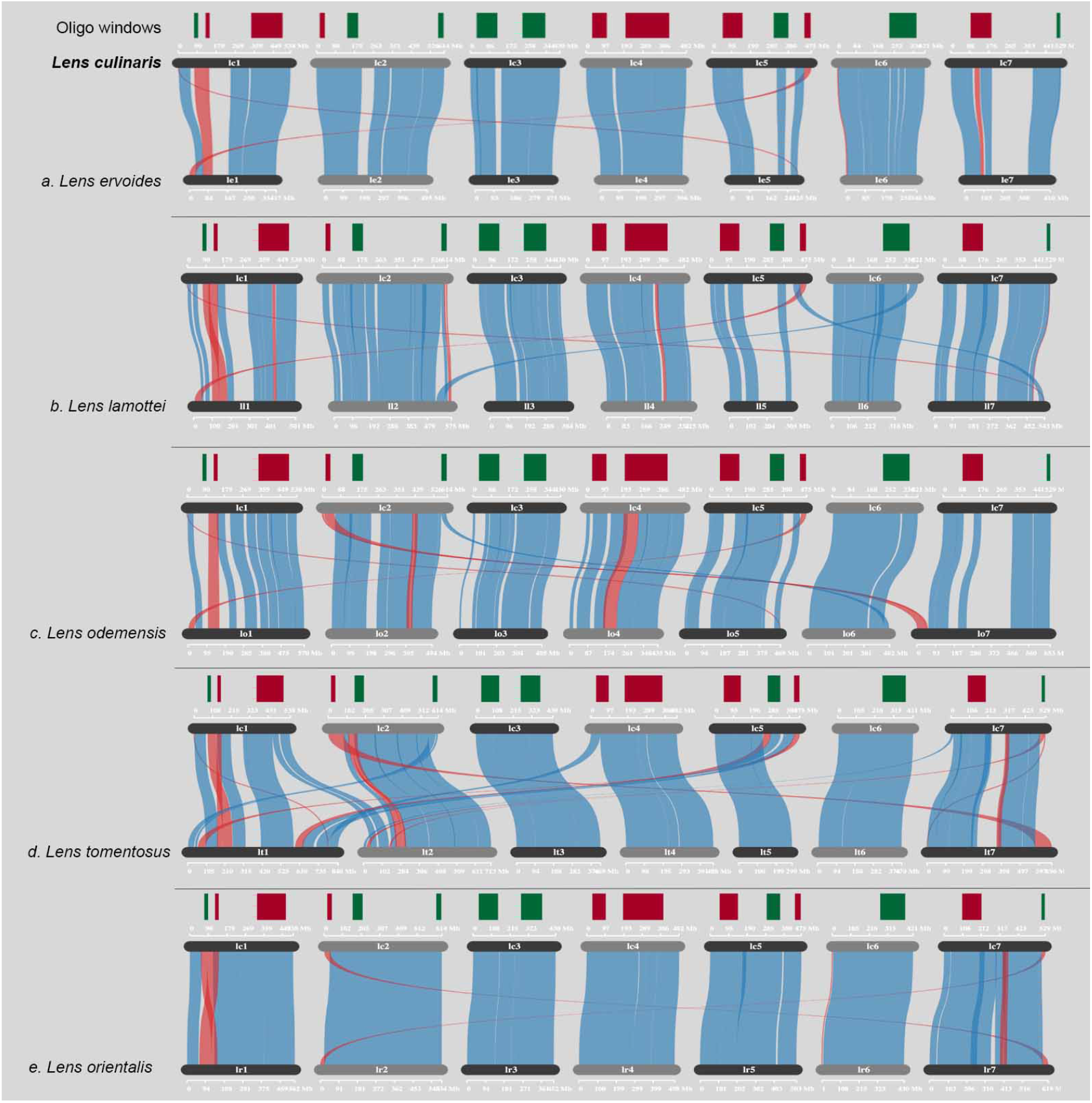
Synteny comparison between Lcu.2RBY chromosomes (Lc#; top in every pair) and those of each of five wild genomes (a. Ler.1DRT, b. Lla.1ESP, c. Lod.1TUR, d. Lto.1BIG, e. Lor.1WPS). Green and red blocks represent the predicted genomic windows in Lcu.2RBY, where probes would cover the maximum number of rearrangements relative to the wild genomes. Blue ribbons represent syntenic genomic regions, and red ribbons represent inverted genomic regions.

### Oligo-FISH probe predictions in *Lens* species

Oligo sequences (green and red) were pre-selected in the chosen genomic windows on the Lcu.2RBY assembly (Table 2). BLASTN analysis confirmed the presence of Lcu.2RBY homologous oligos in five wild genomes - Lor.1WPS, Lto.1BIG, Lod.1TUR, Lla.1ESP and Ler.1DRT - allowing the filtering of probes that should work in a cross-species experiment (Sup. Table 4 and Table 3). Lni.1VIC and Lcu.1GRN were incorporated into the analysis after the initial BLASTN filtering, so they were not included in the Oligo-FISH design.

**Table 3.** Lcu.2RBY oligo probes final coordinates and genomic information.

| Chromosome | Oligo probe | Start | End | Region Length | Number of oligos | Oligos/Kbp | Oligo pool |
| --- | --- | --- | --- | --- | --- | --- | --- |
| Lcu.2RBY.Chr1 | gol11 | 76893900 | 78542425 | 1648525 | 3375 | 2.05 | Lcu.2RBY_Gpool |
| Lcu.2RBY.Chr2 | gol21 | 172423540 | 173547885 | 1124345 | 3375 | 3.00 |  |
| Lcu.2RBY.Chr2 | gol22 | 604707040 | 605269325 | 562285 | 3375 | 6.00 |  |
| Lcu.2RBY.Chr3 | gol31 | 32078260 | 33116187 | 1037927 | 3375 | 3.25 |  |
| Lcu.2RBY.Chr3 | gol32 | 300905060 | 301516885 | 611825 | 3375 | 5.52 |  |
| Lcu.2RBY.Chr5 | gol51 | 348012280 | 348539185 | 526905 | 3375 | 6.41 |  |
| Lcu.2RBY.Chr6 | gol61 | 379187480 | 379713325 | 525845 | 3375 | 6.42 |  |
| Lcu.2RBY.Chr7 | gol71 | 522206540 | 522577327 | 370787 | 3375 | 9.10 |  |
| Lcu.2RBY.Chr1 | rol11 | 147086280 | 149067785 | 1981505 | 3375 | 1.7 | Lcu.2RBY_Rpool |
| Lcu.2RBY.Chr1 | rol12 | 355614280 | 356253425 | 639145 | 3375 | 5.28 |  |
| Lcu.2RBY.Chr2 | rol21 | 36278340 | 36837580 | 559240 | 3375 | 6.03 |  |
| Lcu.2RBY.Chr4 | rol41 | 6279460 | 6765987 | 486527 | 3375 | 6.94 |  |
| Lcu.2RBY.Chr4 | rol42 | 304057720 | 304666605 | 608885 | 3375 | 5.54 |  |
| Lcu.2RBY.Chr5 | rol51 | 57947160 | 58665547 | 718387 | 3375 | 4.7 |  |
| Lcu.2RBY.Chr5 | rol52 | 468283260 | 468760065 | 476805 | 3375 | 7.08 |  |
| Lcu.2RBY.Chr7 | rol71 | 160382620 | 161597887 | 1215267 | 3375 | 2.78 |  |

**Table 4.** Oligo-FISH signal numbers in each *Lens* species’ chromosomes. Species are organized by phylogeny order. Gene pool classification of *Lens* species by Wong et al.,2015.

| Species | Gene Pool | Chr1 | Chr2 | Chr3 | Chr4 | Chr5 | Chr6 | Chr7 | *n*<br>*total* |
| --- | --- | --- | --- | --- | --- | --- | --- | --- | --- |
| *L. culinaris* | 1 | 4 | 3 | 2 | 2 | 2 | 1 | 2 | 16 |
| *L. orientalis* | 1 | 4 | 4 | 2 | 2 | 2 | 1 | 1 | 16 |
| *L. tomentosus* | 1 | 8 | 2 | 2 | 1 | 1 | 1 | 2 | 17 |
| *L. odemensis* | 2 | 3 | 1 | 2 | 2 | 1 | 2 | 3 | 14 |
| *L. lamottei* | 2 | 3 | 2 | 2 | 2 | 2 | 0 | 2 | 13 |
| *L. ervoides* | 3 | 3 | 3 | 2 | 2 | 0 | 1 | 3 | 14 |
| *L. nigricans* | 4 | - | - | - | - | - | - | - | - |

Both Lcu.2RBY_Gpool and Lcu.2RBY_Rpool oligo pools are described in Table 3. Each Lcu.2RBY oligo pool contains 27,000 oligos, derived from 16 regions on the seven *L. culinaris* chromosomes. Based on Lcu.2RBY predictions, the probes should present 16 distinct FISH signals with 3375 oligos each. Gene annotation analysis showed that 16.98% of the oligos are within Lcu.2RBY genes, while the remaining are likely in intergenic regions.

As expected, based on the oligo probe design strategy, most Lcu.2RBY probes have a homologous BLASTN hit within the wild *Lens* genomes and Lcu.1GRN – every BLASTN hit was included as a predicted probe site. The average identity of both Lcu.2RBY oligo pools ranges from 96.21% for *Lens nigricans* genome to 99.79% for *Lens tomentosus* genome, with the highest number of mismatches for *L. nigricans* and the lowest for *Lens orientalis* genomes (Sup. Table 4 and Figure 2). The combination of the rearranged oligos (Figure 2 black dashed lines) and the syntenic oligos (Figure 2 coloured dashed lines) between the cultivated and the wild species provides a barcode for each chromosome from each of the analyzed species (Sup. Figure 1).

**Figure 2.**
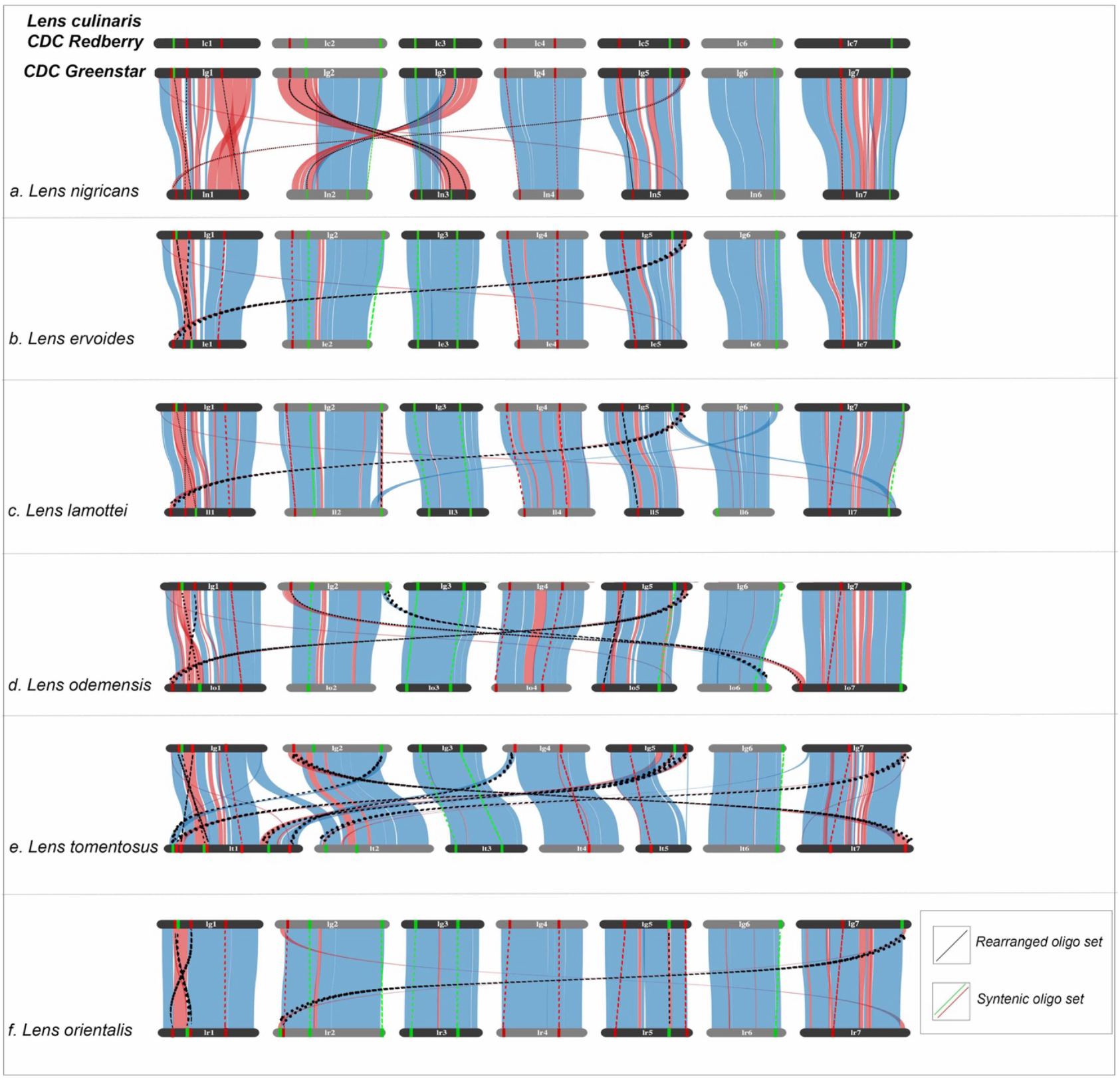
Oligo probes distribution according to BLASTN results in combinations of wild *Lens* genomes (Lni.1VIC, Ler.1DRT, Lla.1ESP, Lor.1WPS, Lod.1TUR, Lto.1BIG) relative to *Lens culinaris* CDC Greenstar (Lcu.1GRN). Lcu.1GRN is being used instead of Lcu.2RBY after quality assessment from Sup. Figure 3. Black dashed lines highlight predicted rearrangements to be confirmed in oligo-FISH. Coloured dashed lines highlight syntenic oligo-FISH signals. Probe signals without dashed lines do not have homologous BLASTN hits.

### Oligo-FISH can be used as a tool to evaluate genome assembly completeness: *Lens culinaris* CDC Redberry vs. CDC Greenstar

The predicted oligo distribution pattern for *L. culinaris* CDC Redberry (Figure 3A) matches the oligo-FISH results for chromosomes 2, 3, 4, 6, and 7. CDC Redberry chromosome 5 (*in situ*) is missing one red signal (rol51) compared to Lcu.2RBY.Chr5 (assembly prediction), which is explained by a longer probe region and, consequently, a lower density of oligos for rol51 (Table 3). Rol51 does not have a high enough probe density to be visible in the FISH in comparison to other oligo probes on chromosome 5.

**Figure 3.**
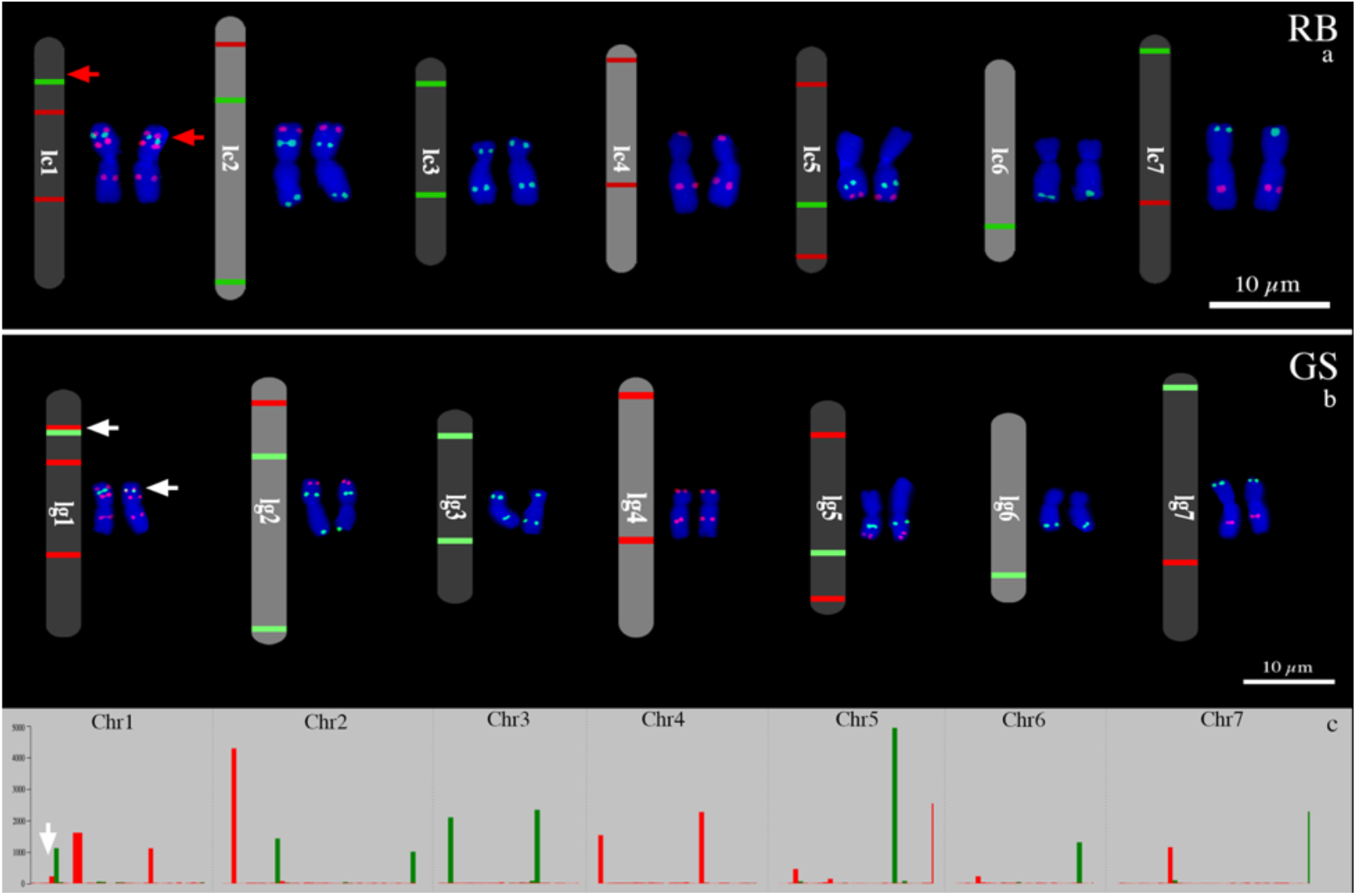
Oligo-FISH results for *Lens culinaris* a) CDC Redberry (RB) and b) Greenstar (GS). c) Oligo density for CDC Greenstar based on BLASTN results. Red arrows point to a missing and an extra red signal on Lcu.2RBY assembly and *in situ* chromosomes, respectively. White arrows pointing to homologous red signals both in Lcu.1GRN assembly and *in situ* chromosomes.

*L. culinaris* CDC Redberry chromosome 1 had an extra red signal when compared to the Lcu.2RBY.Chr1 assembly (Figure 3A, red arrows) that was not detected during probe development. *L. culinaris* CDC Greenstar – another genotype of the same species – had the missing extra red signal both on *in situ* chromosome 1 and in the Lcu.1GRN.Chr1 assembly (Figure 3B and C, white arrows). The extra red signal on CDC Greenstar chromosome 1 belongs to the first red oligo set present in Lcu.2RBY.Chr1 (rol11) (Table 3), which split into two different genomic regions and two distinct oligo-FISH signals in CDC Greenstar (Sup. Figure 2B, black arrow). *L. culinaris* CDC Greenstar genome assembly, therefore, better reflects the *L. culinaris* chromosomes, with a more complete genome assembly. This is backed up by increased BUSCO scores for the Lcu.1GRN assembly (Sup. Figure 3). Because of this, Lcu.1GRN probe distribution was used as a reference for further comparison with the wild *Lens* genomes (Figure 2).

### Genome assembly and synteny confirmations: Oligo-FISH cross-species experiments

Lcu.2RBY oligo probes used for fluorescent *in situ* hybridization (FISH) were successful in six wild *Lens* genomes (Figure 5 and Sup. Figure 4). For most species, the predicted oligo probe locations match the oligo-FISH signals, agreeing with the synteny comparisons in the wild genome assemblies. The *L. orientalis* (Lor) oligo-FISH signal pattern is similar to that of *L. culinaris* CDC Greenstar and CDC Redberry (Lcu) (Sup. Figure 4A). BGE 016880 (Lor) chromosome 1 possesses the homologous extra split red signal present in Lcu oligo-FISH (rol11, see Table 3), which confirms a major inversion on chromosome 1 of Lor relative to Lcu due to its closer localization to the green signal gol11. The results also confirm a chromosomes 2-7 translocation in BGE 016880 (Lor) relative to Lcu.

All *L. tomentosus* (Lto) chromosomes match the genome assembly predictions, confirming the proper chromosome assignment of scaffolds during the assembly process (Figure 4). Lto has the most divergent probe pattern when compared to *L. culinaris,* with multiple translocations and inversions. These results confirm the high number of rearrangements predicted by the synteny analysis, giving assurance that the assembly is correct. Lto chromosome 1 has the extra rol11 signal seen in Lcu.1GRN. The smaller size of Lto chromosome 5 makes the low-density red probe rol51 more visible (see Table 3) (Figure 4B), whereas it is not visible in Lcu or Lor.

**Figure 4.**
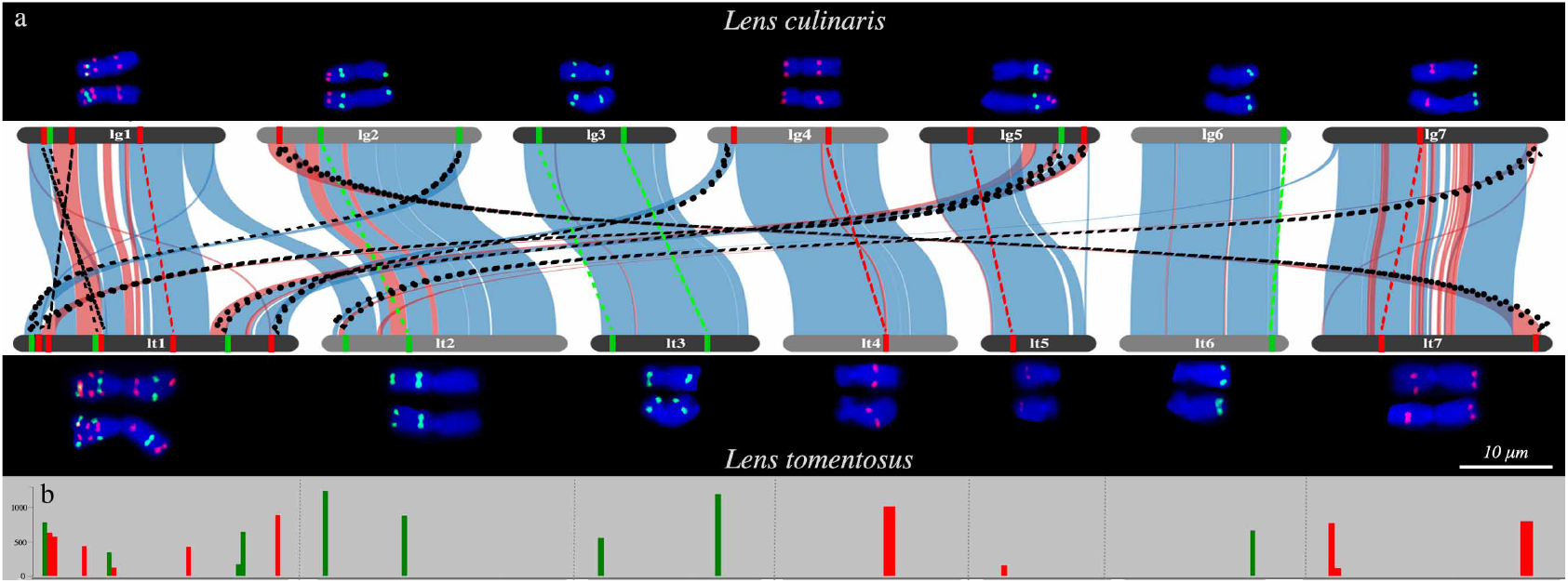
Oligo-FISH results compared to oligo probe predictions for *Lens culinaris* and *Lens tomentosus*. a. Oligo-FISH results alongside oligo probe predictions. b. Oligo density.

Most *Lens odemensis* (Lod) chromosomes match the predicted oligo probe pattern, confirming its genome assembly and synteny analysis. One exception is on chromosome 1, which does not have the extra split red signal seen in Lcu.1GRN (rol11). Chromosome 5 is also missing the low-density red signal (rol51). Chromosome 7 has one missing and one extra red signal that doesn’t match the BLASTN probe density. Thus, it is advised to review the assembly for chromosome 7 in the Lod.1TUR genome (Sup. Figure 4B).

*Lens lamottei* (Lla) has a similar probe distribution to Lod, with most chromosomes in the assembly confirmed through the oligo-FISH results (Sup. Figure 4C). Chromosome 5 is missing rol51 in the low-density probe region, and it has two extra green signals that are not present in the BLASTN probe density distribution (Sup. Figure 4C.2). Chromosome 6 is missing a green signal, which is present in the opposite location when compared to Lcu.1GRN. The Lla.1ESP assembly chromosomes 5 and 6 should be reviewed.

*Lens ervoides* (Ler) has a similar probe pattern to Lla (Sup. Figure 4D). Once again, chromosome 5 is missing the red signal (rol51) present on the BLASTN due to a low density (Table 3). In the same chromosome, there is no sequence homologous to the green signal gol51 that is found in Lcu.1GRN, likely lost during species differentiation since Ler is distantly related to Lcu. *L. nigricans* (Lni) is the most distantly related species when compared to the reference Lcu and is also the only species in which the oligo probes did not produce a clear pattern of signals (Sup. Figure 5A) despite the assembly (Lni.1VIC) having been predicted to have high oligo density (Sup. Figure 5B). When analyzing the distribution of these oligos for each Lcu.1VIC chromosome, there was a noticeably higher dispersion of probes, as compared to in the other *Lens* genomes, creating more non-specific signals, as seen in the oligo-FISH results. Oligos tend to be less conserved in distantly related species (Braz et al., 2018; Braz, Yu, et al., 2020), reducing the likelihood of producing sufficient signals for microscope capturing.

### Precise *Lens* karyotyping and chromosome evolution

The oligo probe patterns allow us to precisely define the homologous chromosomes in six of the seven analyzed *Lens* species, building a strong karyotype system (Figure 5). *L. culinaris* and *L. orientalis* possess a similar karyotype. Both species have 16 oligo-FISH signals on their chromosomes (Table 4), highlighting their sequence homology, while the arrangement of those probes reflects their few structural differences, notably chromosomes 2 and 7. *L. tomentosus* also has a similar number of oligo-FISH signals compared to Lcu and Lor, with a single addition of the rol51. Those results attest to the higher level of sequence homology among the closest related *Lens* species from the primary gene pool (Table 4).

Despite the close phylogenetic relatedness of *L. tomentosus* to both Lcu and Lor, this is not obvious when using Lto oligo-FISH signal arrangements, as they differ drastically from the other two species in chromosomes 1, 2, 4, 5 and 7 due to multiple rearrangements Lto chromosome 1, the most rearranged chromosome relative to all other *Lens* spp., has double the number of oligo-FISH signals when compared to Lcu and Lor chromosomes 1 (Figure 5, Table 4). This concentration of probes in one chromosome highlights how affected the remaining chromosomes of Lto are due to the genomic changes. For example, chromosome 5, is visibly smaller in the species compared to others.

**Figure 5.**
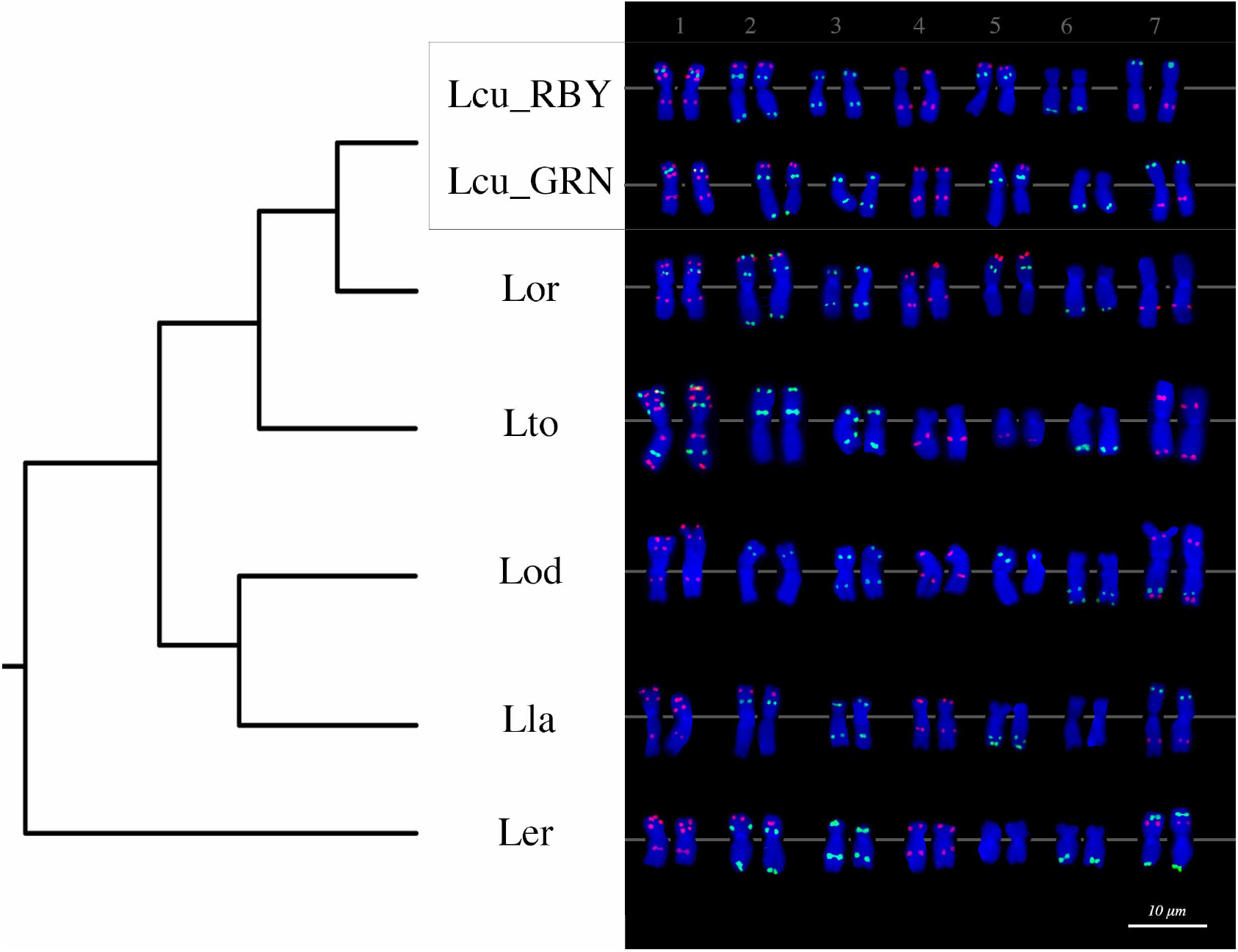
*Lens* species karyotype based on Oligo-FISH. Species codes proposed as Table 1: Lcu: *L. culinaris*, Lor: *L. orientalis*, Lto: *L. tomentosus*, Lod: *L. odemensis*, Lla: *L. lamottei*, Ler: *L. ervoides*. Phylogeny proposed by Wong et al., 2015.

The reduction of oligo-FISH signals according to *Lens* species differentiation (Table 4) is related to the oligo sequence divergence from *L. culinaris* – the oligo reference –to the more diverged species such as Lod, Lla and Ler (Braz et al., 2018; Braz, Yu, et al., 2020; do Vale Martins et al., 2021). *L. odemensis* has a similar karyotype in probe signal, number and arrangement to *L. lamottei* (Figure 5, Table 4). The similar probe signal number reflects a higher sequence homology among these two species rather than with species from the primary gene pool (Lcu, Lor and Lto). Lod and Lla differ in chromosomes 2, 5, 6 and 7 by five oligo signals (Figure 5, Table 4). *L. ervoides* also has a similar karyotype to Lod and Lla, with 14 oligo-FISH signals, even though the species is evolutionarily more diverged. Chromosomes 1, 3 and 4 are identical in oligo-FISH signals among Ler, Lod and Lla.

### Oligo-FISH probes can be used to detect intraspecies structural variation in *Lens* species

*L. orientalis*, the closest relative to *L. culinaris,* is often used in plant breeding programs for the introgression of beneficial alleles from the wild parent (Singh et al., 2018). The Lcu.2RBY oligo probes were applied to a second accession of *L. orientalis,* IG 72643, and compared with BGE 016880 – the first *L. orientalis* accession used in this study (Figure 6).

**Figure 6.**
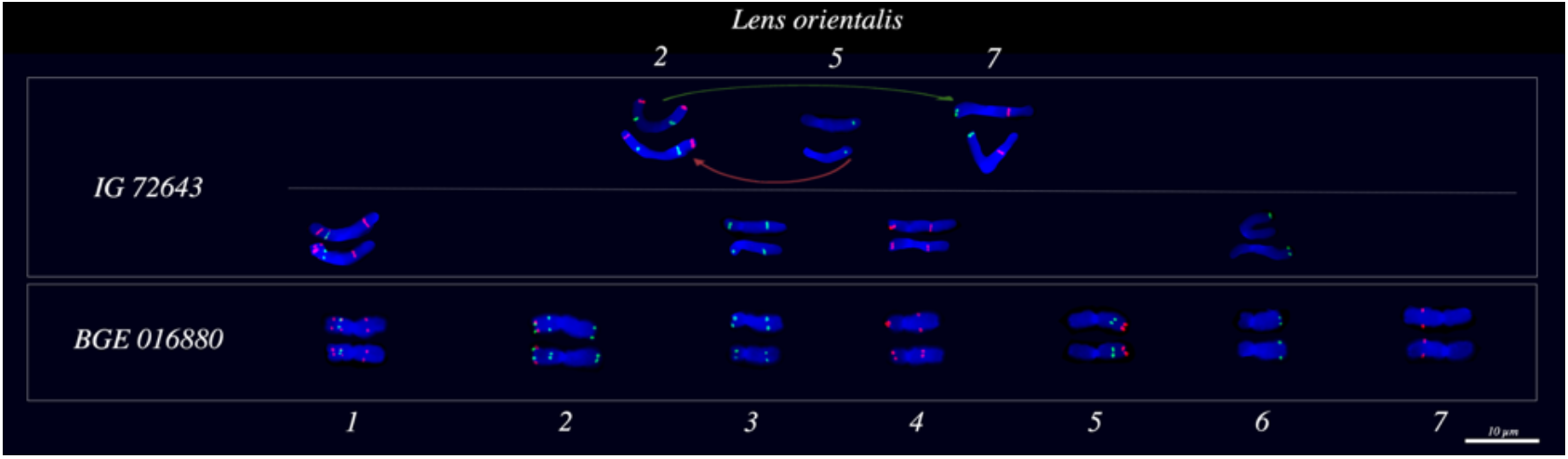
Oligo-FISH analysis of *Lens orientalis* IG 72643 in comparison to BGE 016880. Green arrow points to a proposed translocation between chromosomes 2 and 7. Red arrow points to a proposed translocation between chromosomes 5 and 2.

Chromosomes 1, 3, 4 and 6 of IG 72643 have identical probe distribution when compared to BGE 016880, while chromosomes 2, 5 and 7 differ between the two accessions. Chromosome 2 has one extra red signal and a missing green signal in IG 72643, chromosome 5 is missing one red signal, and chromosome 7 has an extra red signal. The arrangement of those probes is different only in IG 72643 chromosome 2.

It is possible to propose rearrangements that led to the karyotype differentiation between these two accessions. A translocation between chromosomes 2 and 7 could lead to the extra green signal in 7 (Figure 6, green arrow), while another translocation between chromosomes 5 and 2 could lead to the extra red signal in 2 (Figure 6, red arrow). Those rearrangements will be further confirmed after finalizing the genome assembly of IG 72643 and performing synteny analysis with BGE 016680.

## Discussion

Oligo-FISH studies across multiple species primarily rely on the genome assembly of a single species, with the probes then utilized in species lacking a genme assembly to detect structural variation (Braz et al., 2018; do Vale Martins et al., 2021; Jiang, 2019; G. Li et al., 2021). We developed an oligo-FISH probe system for karyotyping across the genus *Lens,* starting with the cultivated *L. culinaris* genome assembly (Lcu.2RBY) (Table 3), followed by synteny analysis and homology filtering against the genome assemblies of five *Lens* spp. wild relatives (Figures 1 and 2).

Advancements in genome sequencing technologies are supporting the release of high-quality genome assemblies; however, limitations still exist, primarily related to the assembly of large, highly repetitive genomes, such as those of *Lens* spp. Ultra-long-read sequencing technologies and contact mapping have helped, but it still requires a large investment in sequencing to achieve the required depth to be confident in an assembly. The accuracy of synteny analysis can be affected when comparing highly rearranged genomes like those of *Lens* spp., among themselves. The use of various strategies, such as cytogenetic maps, for validating the genome assembly and synteny analysis is thus highly recommended (Li & Durbin, 2024; Luo et al., 2021).

Our *Lens* oligo-FISH system innovatively enables the validation of proper genome assembly scaffolding into chromosomes and confirms the proposed synteny among *Lens* species. Oligo-probe-based genomic predictions match most of the *Lens* physical chromosomes, attesting to the quality of the genome assembly scaffolding process, even using highly repetitive genomes. The changes in most probe positions for *L. culinaris* and its wild relatives confirm the rearrangements detected in the synteny analysis (Figure 5A and Supplementary Figure 4) (Silvestrini et al. Unpublished). The probes are also useful for suggesting improvements to the assembly of specific chromosomes, such as Lod.1TUR chromosome 7, where the predictions do not match the oligo-FISH signals due to misassembled regions (Sup. Figure 4B).

Differences in the predictions of oligo probes for Lcu.1GRN and Lcu.2RBY that were then confirmed with the oligo-FISH could be used as a quality assessment, confirming the missing signal on CDC Redberry chromosome 1 (Figure 3) was related to the assembly which is also borne out by the higher fragmented BUSCO scores (Sup. Figure 3). Lou et al., 2014 reported the use of oligo-FISH painting probes to assess genome assembly completeness in *Cucumis* species using meiotic pachytene cells. This, however, is the first time that single-locus oligo probes in mitosis have been reported for assessing genome assembly quality.

The oligo probes enable us to propose the most precise karyotype for six *Lens* species to date, accurately defining each pair of chromosomes without the need for chromosome measurements or the use of repetitive DNA FISH probes. Diversity in karyotypic formulas and differences in satellite DNA FISH probes, due to the repetitive nature of *Lens* genomes (Silvestrini et al., Unpublished), have been proposed for the seven *Lens* species suggesting multiple rearrangements (Balyan et al., 2002; Jha, 2022; Fernández et al., 2005; Gaffarzade et al., 2007; Galasso, 2003; Jha & Halder, 2016; Ladizinsky & Abbo, 1993; Salimuddin & Ramesh, 1994). These have now been confirmed physically and precisely in our work. These rearrangements altered the structure of *Lens* species chromosomes, as determined by oligo-FISH hybridization patterns, throughout evolution, without changing their sequence composition, thereby reflecting the phylogenetic proximity of these species (Alo et al., 2011). We also confirm the limitation of oligo-FISH probes when conducting experiments with more distantly related species (Braz et al., 2018; Braz, Yu, et al., 2020; Hoang et al., 2021), with the lack of clear signals on *L. nigricans* chromosomes (Figure 4). We detected intraspecific variation in two different genotypes of *L. orientalis*. The occurrence of distinct structural rearrangements within this species has already been proposed (Ladizinsky, 1979; Ladizinsky et al., 1984b, 1985; Ladizinsky & Abbo, 1993) based on examination of chromosome pairing during meiosis.

The oligo-FISH probes are useful for characterizing and detecting structural variation among genotypes of the same or different *Lens* species without the need for a genome assembly. This would allow researchers to screen wild relatives not just for phenotypes of interest but also allow them to pick accessions that will more likely lead to successful pairing and recombination during F1 meiosis. Crossings between species from different gene pools can bring undesirable alleles into the cultivated *L. culinaris* genome due to linkage drag associated with the numerous rearrangements. The probes are useful for screening populations and tracking specific chromosome combinations, allowing breeders to stack desirable alleles while overcoming linkage drag issues. Chromosome 7 is a good example of the rearrangement’s imposition when trying to transfer alleles, considering it is a chromosome known to be more prone to rearrangement than others (Ramsay et al., 2021; Silvestrini et al., Unpublished).

The developed oligo-FISH probe system in *Lens* spp. is useful for assessing genome assembly quality and completeness, as well as for further screening of populations and intra/interspecific hybrids in plant breeding programs. Chromosome behaviour during reproduction, facing larger structural variation, is also of interest for crop production and genome evolution studies. Our cross-species oligo-FISH system enables future studies in the entire *Lens* genus regarding chromosome pairing, recombination and hybrid viability for the seven species. The probes are also to be tested in other legume species, allowing even further studies in legume genomics. This is a remarkable achievement for legume science and breeding, with an oligo-FISH system that sheds light on the hidden structural variation of those big and repetitive-rich genomes.

## Data availability

Genome assemblies are available at https://knowpulse.usask.ca/genome-assembly/insert specific genome assembly code from Table 1.

Oligo sequence information is available at https://knowpulse.usask.ca/Analysis/Oligo-FISH-probe-design-for-Lens-rearrangements. Sequences used to develop Lcu.2RBY oligo pools are available in the Supplementary Tables file.

## Acknowledgments

We thank Arbor Biosciences scientists for their support with the Oligo probes development and Jiri Macas’ group from The Czech Academy of Sciences for providing support for the slide preparation protocol.

This work was supported by the ‘Enhancing the Value of Lentil Variation for Ecosystem Survival (EVOLVES)’ project funded by Genome Canada [grant: LSP18-16302] and managed by Genome Prairie. Matching funding was provided by: Saskatchewan Pulse Growers [grant: BRE 1516], Western Grains Research Foundation [grant: GC1903], Saskatchewan Ministry of Agriculture [grant: 20200026], BASF, and the University of Saskatchewan.

