## Supplementary Figures for "Oligo-FISH Validates Genome Assemblies and Delivers the Most Precise Karyotype for *Lens* Mill. Species"

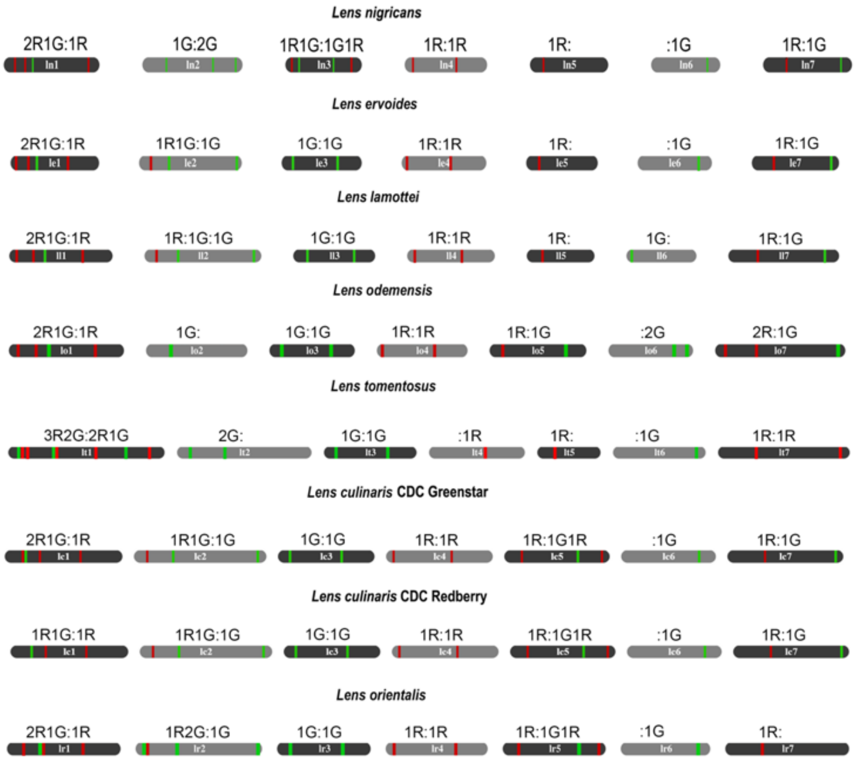


Supplementary Figure 1. Oligo barcode system based on BlastN results for each *Lens* species chromosome.

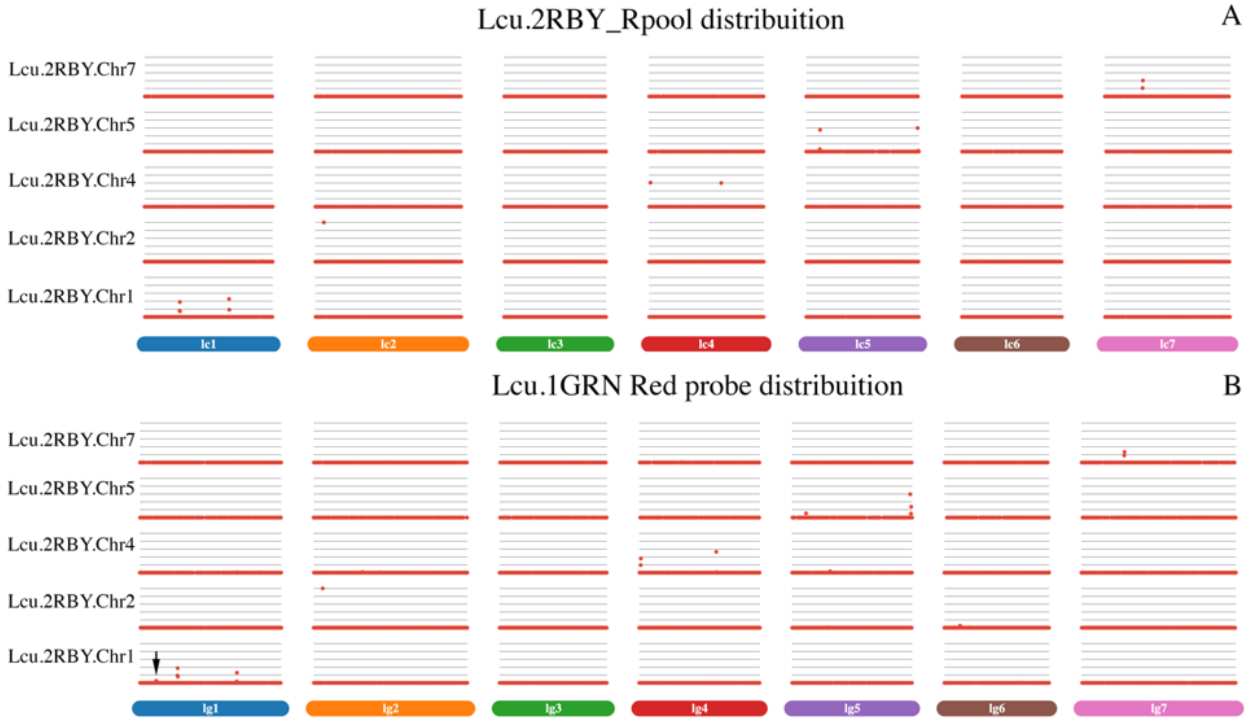


Supplementary Figure 2. Lcu.2RBY and Lcu.1GRN red probe distribution. The black arrow highlights the origin of the extra red signal on Lcu.1GRN.Chr1 as Lcu.2RBY.Chr1. X-axis: Lcu.2RBY and Lcu.1GRN genome assemblies. Y-axis: Lcu.2RBY_Rpool distribution.

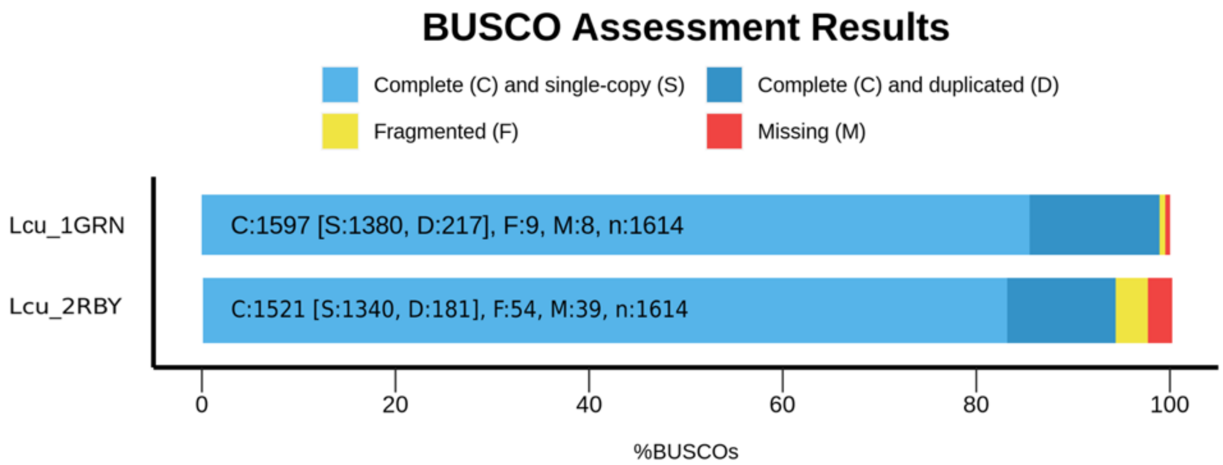


Supplementary Figure 3. BUSCO Scores for assessing genome completeness for Lcu.1GRN and Lcu.2RBY.

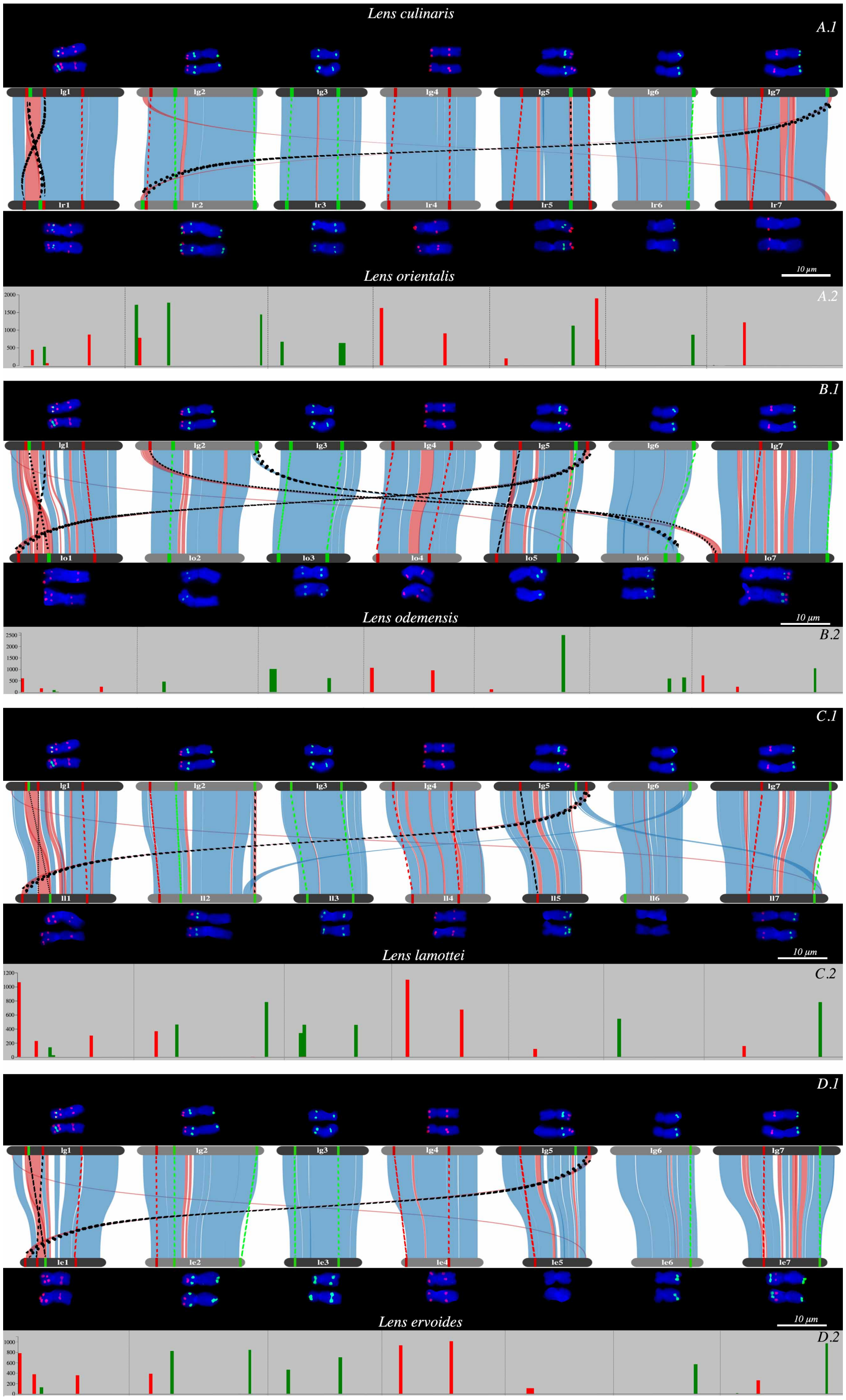


Supplementary Figure 4. Oligo-FISH results compared to oligo probe predictions for wild *Lens* species. .1 *Lens culinaris* CDC Greenstar oligo-FISH and probe prediction compared to each wild species oligo-FISH and probe prediction. Oligo-FISH and oligo probe predictions. .2 Oligo density based on BlastN results.

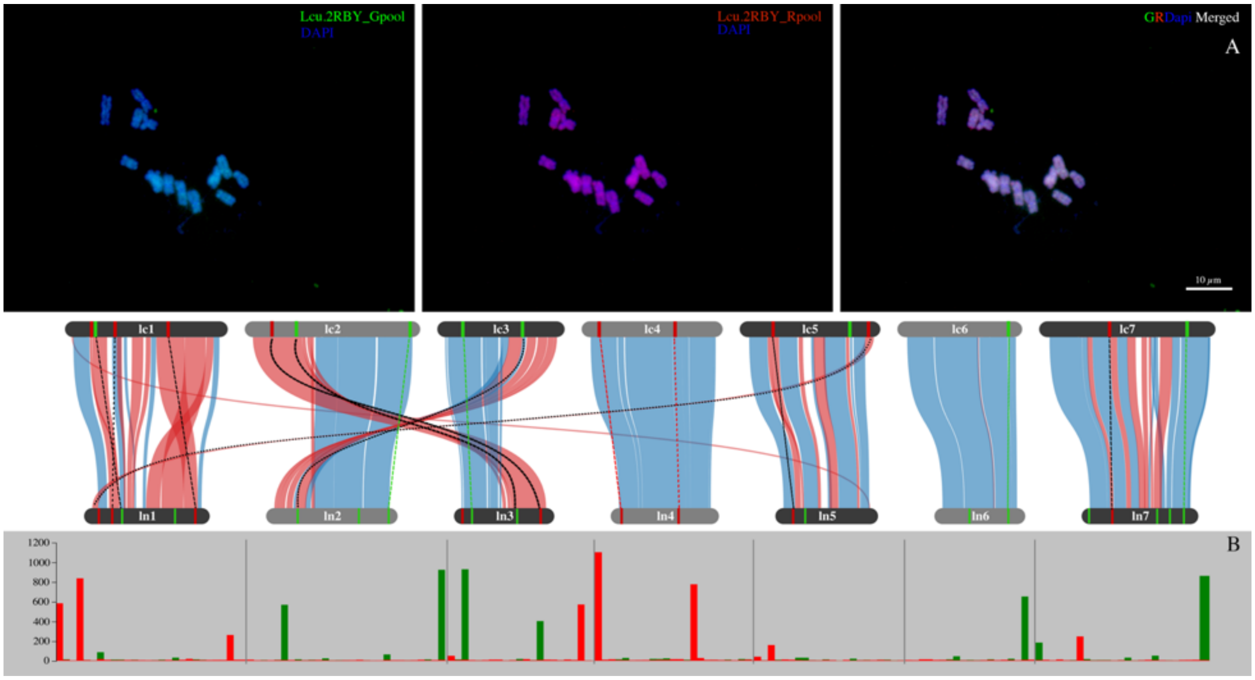


Supplementary Figure 5. *Lens nigricans* Oligo-FISH highlight the lack of a clear pattern of signals for both Lcu.2RBY oligo probes. A: Oligo-FISH images. B: BlastN probe density for Lni.1VIC chromosomes.
